# Comparative evidence for the independent evolution of hair and sweat gland traits in primates

**DOI:** 10.1101/430454

**Authors:** Yana G. Kamberov, Samantha M. Guhan, Alessandra DeMarchis, Judy Jiang, Sara Sherwood Wright, Bruce A. Morgan, Pardis C. Sabeti, Clifford J. Tabin, Daniel E. Lieberman

**Affiliations:** Department of Genetics; Harvard Medical School; Boston, MA 02115; USA; Department of Human Evolutionary Biology; Harvard University; Cambridge, MA, 02138; USA; Department of Dermatology, Cutaneous Biology Research Center; Massachusetts General Hospital and Harvard Medical School; Charlestown, MA, 02129; USA; FAS Center for Systems Biology, Department of Organismic and Evolutionary Biology; Harvard University; Cambridge, MA, 02138; USA; The Broad Institute of MIT and Harvard; Cambridge, MA, 02142; USA; Howard Hughes Medical Institute; Chevy Chase, MD, 20815; USA; Department of Immunology and Infectious Disease, Harvard T.H. Chan School of Public Health, Boston, MA, 02115; USA

**Keywords:** Skin, Human evolution, Sweat gland, Hair, Eccrine gland, Ectodermal appendage

## Abstract

Humans differ in many respects from other primates, but perhaps no derived human feature is more striking than our naked skin. Long purported to be adaptive, humans’ unique external appearance is characterized by changes in both the patterning of hair follicles and eccrine sweat glands, producing decreased hair cover and increased sweat gland density. Despite the conspicuousness of these features and their potential evolutionary importance, there is a lack of clarity regarding how they evolved within the primate lineage. We thus collected and quantified the density of hair follicles and eccrine sweat glands from five regions of the skin in three species of primates: macaque, chimpanzee and human. Although human hair cover is greatly attenuated relative to that of our close relatives, we find that humans have a chimpanzee-like hair density that is significantly lower than that of macaques. In contrast, eccrine gland density is on average 10-fold higher in humans compared to chimpanzees and macaques, whose density is strikingly similar. Our findings suggest that a decrease in hair density in the ancestors of humans and apes was followed by an increase in eccrine gland density and a reduction in fur cover in humans. This work answers longstanding questions about the traits that make human skin unique and substantiates a model in which the evolution of expanded eccrine gland density was exclusive to the human lineage.

## 1. Introduction

Human skin is predominantly populated by a mixture of hair follicles and water-secreting eccrine sweat glands (Szabo, 1967; Montagna and Parakhal, 1974) both of which derive from the embryonic ectoderm (Biggs and Mikkola, 2014). These ectodermal appendages are critical components of our species’ main mechanism of cooling: the evaporation of water secreted by eccrine sweat glands from the skin surface (Montagna and Parakhal, 1974). Thermoregulatory sweating occurs in great apes and Old World monkeys (Johnson and Elizondo, 1974, 1979; Hiley, 1976; Whitford, 1976; Elizondo, 1977; Kolka and Elizondo, 1983), however, humans are exceptional among catarrhines in their dramatically enhanced sweating capabilities, which are exclusively dependent on the activity of eccrine glands (Folk, 1974; Hiley, 1976; Folk and Semken, 1991). Because a loss of fur cover leads to increased rates of convection and evaporation (Hardy, 1953; Dale et al., 1967; Wheeler, 1991), it has been proposed that the apparent nakedness of human skin enhances the effectiveness of eccrine sweating as a cooling mechanism (Carrier et al., 1984; Wheeler, 1984, 1991; Lieberman, 2015). However, the developmental origins of this striking appearance are unclear. Hypotheses to explain the human condition include overall reduction in hair density (Schultz, 1931; Newman, 1970; Schwartz and Rosenblum, 1981; Wheeler, 1984; Folk and Semken, 1991; Sandel, 2013), miniaturization of body hair (Montagna, 1963, 1985; Montagna et al., 1992), or both (Schwartz and Rosenblum, 1981; Sandel, 2013). To our knowledge, the only previously published study of comparative hair density in humans and other primates evaluated only hairs larger than 2mm in length (Schultz, 1931). This cut-off excludes the microscopic vellus hairs that dominate human skin (Szabo, 1967) thus drastically under-reporting human hair density (Montagna, 1963; Montagna and Parakhal, 1974). More recent analyses of human body hair report increased estimates of follicular densities, however no other species were evaluated in these studies (Montagna et al., 1962; Szabo, 1967) making it difficult to draw conclusions of comparative hair composition.

The human thermoregulatory skin apparatus has been proposed to rely not only on adaptive changes in follicular make-up, but also on a high density of eccrine sweat glands in the hairy skin (Montagna, 1963, 1972; Newman, 1970; Carrier, 1984; Wheeler, 1984; Folk and Semken, 1991; Bramble and Lieberman, 2004; Lieberman, 2015). An innovation of mammals, eccrine glands appear to have first evolved in the volar skin to enhance traction (Adelman et al., 1975). However, their distribution was expanded into the hairy skin in the common ancestor of catarrhines, the group of primates that includes Old World monkeys (Montagna et al., 1964, 1966; Montagna, 1972), great apes (Ellis and Montagna, 1962; Montagna and Yun, 1963; Montagna, 1972) and humans (Montagna, 1963; Szabo, 1967). While it remains the subject of debate why an expanded distribution of eccrine glands initially evolved and was retained in higher primates (Montagna, 1972; Schwartz and Rosenblum, 1981; Wheeler, 1984; Folk and Semken, 1991; Sandel, 2013; Best and Kamilar, 2018), in humans these organs are indispensable for heat dissipation (Kuno, 1956; Montagna, 1972). Humans are often claimed to have the highest density of eccrine glands of any primate, with great apes having an intermediate density and Old World monkeys having the lowest (Montagna, 1963, 1972; Newman, 1970; Montagna and Parakhal, 1974; Folk and Semken, 1991). These widely accepted assertions are based on limited data. There are very few studies on the density of anatomical eccrine glands in the hairy skin of humans (Wagner, 1844; Kuno, 1956; Szabo, 1962). The acquisition of reliable data on human eccrine gland density is challenged by the need to biopsy the skin to accurately score these organs. As a result, most studies of human gland density do not evaluate total anatomically present glands but rather the subset of glands which are active, a trait that is heavily influenced by complex environmental factors such as humidity and temperature during early childhood (Kuno, 1956; Knip, 1977). An additional complication is that comparative eccrine gland estimates from non-human primates have largely been reported as ordinal values relative to the abundance of apocrine glands, another type of ectodermal appendage that is not found in most human skin, rather than as explicit densities (Ellis and Montagna, 1962; Montagna and Yun, 1963; Montagna et al., 1964, 1966; Montagna, 1972). A study of sweating in chimpanzees demonstrates that apocrine glands are activated in response to thermal challenge and are major contributors to heat loss from the skin in this species (Whitford, 1976). Thus, published estimates of eccrine gland density in non-human primates from studies of water and heat loss in these species (Johnson and Elizondo, 1974, 1979; Whitford, 1976; Kolka and Elizondo, 1983) are confounded by a lack of distinction between the relative contribution of apocrine versus eccrine sweating. Consequently, we lack a reliable quantitative comparison of eccrine gland density across primates calling into question our understanding of the so-called uniqueness of the human eccrine gland phenotype.

To characterize how human ectodermal appendage phenotypes differ from those of our closest primate relatives, we systematically surveyed hair follicle and eccrine gland densities in five body regions (forehead, back, chest, forearm and thigh) in three species of catarrhines: human (*Homo sapiens*), common chimpanzee (*Pan troglodytes*) and rhesus macaque (*Macaca mulatta*). Our results provide a quantitative assessment of the differences in skin appendage composition between macaques, chimpanzees and humans and establish the necessary phenotypic context for pursuing the underlying genetic changes driving the emergence of two long-appreciated thermoregulatory attributes of human skin.

## 2. Methods

### 2.1 Tissue Sources

Full thickness post-mortem skin biopsies from seven humans (4 females, 3 males), four chimpanzees (2 females, 2 males) and eight rhesus macaques (5 females, 3 males) were used in this study (Fig. 1 and SOM Table S1). All individuals were adults, and five body regions that contain eccrine glands were sampled: forehead, back, chest, dorsal thigh and dorsal forearm. Samples from non-human primates were taken post mortem and human tissues were harvested from subjects who had donated their bodies for scientific research. Human skin samples were obtained with permission for the use of cadaveric material from the Anatomical Gift Program at Harvard Medical School from individuals who had donated their bodies for research. Skin samples from designated regions from one female and one male chimpanzee who died of natural causes were obtained with permission from the Yerkes National Primate Research Center. In addition, we collected skin biopsies from two wild-caught adult chimpanzee cadavers which were part of the Museum of Comparative Zoology Collection. Macaque skin samples were obtained from designated body regions from post-mortem biopsies of animals from the macaque colony at the New England Primate Research Center. Individuals used in this study had no known or visible skin pathologies. Information on individuals used in this study is listed in Supplementary Online Material (SOM) Table S1). All experiments were conducted in accordance with the permissions obtained from: the Anatomical Gift Program at Harvard Medical School, the Yerkes National Primate Research Center and the New England Primate Research Center.

**Fig. 1.**
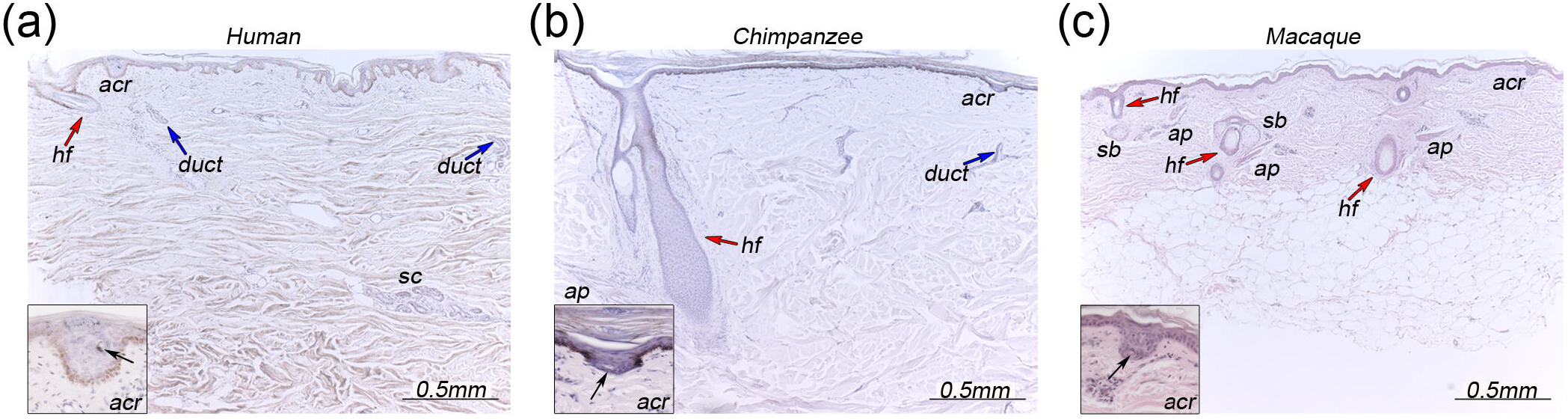
Hair and eccrine glands in the skin of three primates. a – c) Representative images of skin sections taken from the back of human (a), chimpanzee (b) and macaque (c) study subjects. Insets are magnified views of the acrosyringium (acr), the most apical portion of the eccrine gland, showing characteristic coiling of the duct (black arrow- lumen of duct). Secretory coil of eccrine gland (sc), hair follicle (hf) and red arrow, eccrine gland duct (blue arrow), sebaceous gland (sb) and arrector pili muscle associated with hair follicles (ap).

### 2.2 Tissue sampling

For consistency, body areas for sampling were chosen as follows. Forehead skin: lateral, above the brow. Back: inferior angle of the scapula. Thigh: dorsal at 1/2 length. Forearm: dorsal at 1/2 length. Chest: level of sternal angle, midpoint of torso half. Full thickness biopsies of skin from the left and right side of the body were taken from each individual for each body region and used in analysis. All macaque and captive-bred chimpanzee skin samples were collected at necropsy shortly after death.

### 2.3 Tissue processing

All macaque and captive-bred chimpanzee samples were received in cold phosphate buffered saline (PBS), washed 3 times in 100% PBS to ensure full hydration and had images taken of each using a Leica MZFLIII stereomicroscope equipped with a Nikon DXM1200F camera with a scale bar to allow for quantification of area. Embalmed human and wild-caught chimpanzee samples were rehydrated in phosphate buffered saline after collection and images taken of each sample as before. Area was calculated from images with scale bar using ImageJ software. We cannot exclude the possibility that embalming may influence the size of the area detected, however, the densities we obtained are in agreement with those of prior studies on eccrine gland density from fresh human skin samples (Wagner, 1844; Kuno, 1956) and, for chimpanzees, consistent with the densities detected from captive animals in our study, indicate that these effects are likely minimal. Fresh skin samples were fixed in paraformaldehyde and washed in PBS. All skin samples were processed through a series of dehydration steps in ethanol into Xylenes (Thermo-Fisher) and embedded in paraffin for histological sections. All samples were sectioned at 15 microns and slides stained with Hematoxylin (Sigma Aldrich) and Eosin (Sigma Aldrich) according to standard protocols. Post-staining, images of serial sections were taken on a Leica DM5500B microscope equipped with a Leica DEC500 camera.

### 2.4 Quantification of ectodermal appendages

Hair follicles and sweat glands were identified morphologically by their distinct appearance in serial sections from each tissue sample. A hair follicle was counted only if the opening of the hair canal and follicle were visible in the analyzed section. Eccrine glands were scored if duct, acrosyringium and opening onto the skin surface were visible in a section. The number of each appendage type, or total appendage types, was divided by the calculated area of the whole tissue sample to determine the density of appendages. A minimum of two replicates were evaluated from each body region for each individual in our study and the average density was used in comparative analysis. Raw quantification and imaging data for samples used in this study are available upon request to Yana G. Kamberov and Daniel E. Lieberman.

### 2.5 Statistical analysis

Ordinary one-way ANOVA followed by Sidak multiple comparisons test with a single pooled variance was performed on the means for each body region using GraphPad Prism version 7.00 for Windows, GraphPad Software, La Jolla California USA, www.graphpad.com.

## 3. Results

### Study subjects and analysis of anatomical ectodermal appendage densities

To assess hair and eccrine gland densities, we collected post-mortem full thickness skin biopsies from five body regions (forehead, back, chest, dorsal thigh, and dorsal forearm) of macaque, chimpanzee and human subjects (Fig.1). After tissue collection, the surface area of each tissue sample was calculated. The number of hair follicles and eccrine glands in each biopsied sample were assessed across histologically-stained serial sections, and density was calculated (SOM Table S2). Evaluation of calculated densities from biological replicates of each surveyed area did not reveal significant differences in the density of either appendage type. Thus, densities used for comparative analysis were calculated based on the mean value of all replicate samples for each area.

We report the densities of hair follicles and sweat glands for each sample per unit area. Evaluation of densities scaled to body mass did not alter the trends observed in our analysis and we did not observe sex-based differences in either eccrine gland or hair follicle density in any species (data not shown).

### Reduction in hair density distinguishes hominids from macaques but not each other

Analysis of hair density revealed that in most body regions, macaques have approximately two to twenty one fold higher average hair density than either chimpanzees or humans (Fig. 2a – d, Sidak multiple comparisons test adjusted *P* values: Forehead *P* = 0.03 (macaque (M) : chimpanzee (C); *P* = 0.04 (M: human (H)); Back *P* = 0.004 (M:C); *P* < 0.0001 (M:H); Dorsal Thigh *P* = 0.002 (M:C); *P* = 0.0001 (M:H); Dorsal Forearm *P* = 0.0030 (M:C); *P* = 0.0011 (M:H)). In the chest, however, macaques had a higher hair density than humans but not chimpanzees (Fig. 2e, adjusted *P* values: *P* = 0.2160 (M: C); *P* = 0.0039 (M: H)). In contrast, average human and chimpanzee hair densities did not differ significantly in any region, despite the very different external appearances of these species (Fig. 2a – e, adjusted *P* > 0.05 for all areas). We note that only two chimpanzee individuals and four human subjects were available for analysis of appendages in the chest, which may help to explain why this region is an outlier in this and subsequent analyses.

**Fig. 2.**
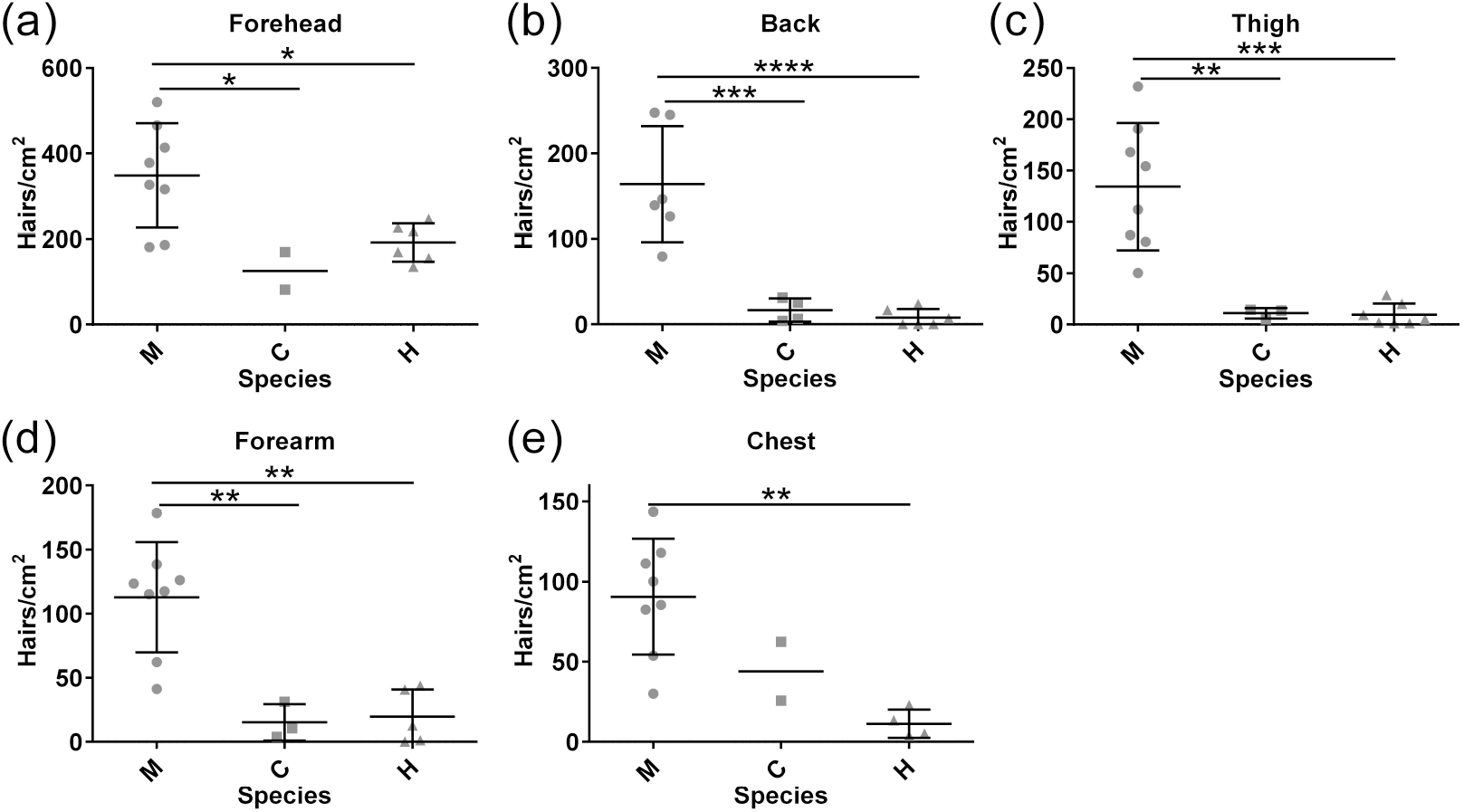
Hair density across body regions in three primates. a – e) Human hair density does not differ significantly from that of chimpanzees, but is decreased relative to macaques. Each point represents the average density of hair follicles in biological replicates for each individual analyzed. Mean and standard deviation are shown for each species. Standard deviation is not shown when sample size did not exceed two individuals. Abbreviations: macaque (M), chimpanzee (C) and human (H). Significance: * *P* < 0.05, ** *P* < 0.01, *** *P* < 0.001, **** *P* < 0.0001.

In humans and macaques, for which we had the largest sample size, significantly greater hair density was detected in the forehead relative to other sampled regions. No significant difference in hair density was observed between the other sampled body regions (SOM Fig. S1a, b and data not shown). We noted a similar trend in chimpanzees (SOM Fig. S1c).

### Increased eccrine gland density distinguishes humans from other surveyed primates

Consistent with previous reports (Montagna, 1963, 1972; Szabo, 1967) eccrine glands were present in all body regions analyzed in the three species (Fig. 3). Measured eccrine gland densities did not vary significantly between chimpanzees and macaques (adjusted *P* > 0.05 for all body regions), however, human eccrine gland density was strikingly higher in virtually all sampled body regions (adjusted *P* < 0.0001 for all regions), averaging ten-fold higher density compared to both macaques and chimpanzees (Fig. 3). In humans, the absolute density of eccrine glands in the forehead was on average 2.5 fold higher (adjusted *P* < 0.0001 for all comparisons) than in the other four sampled areas (SOM Fig. S2). The lowest eccrine gland density detected was in the chest (SOM Fig. S2). In contrast, eccrine gland density was not significantly different between macaques and chimpanzees (adjusted *P* > 0.05 for all comparisons), neither of which had higher densities in the forehead (Fig. 3, SOM Fig. S2).

**Fig. 3.**
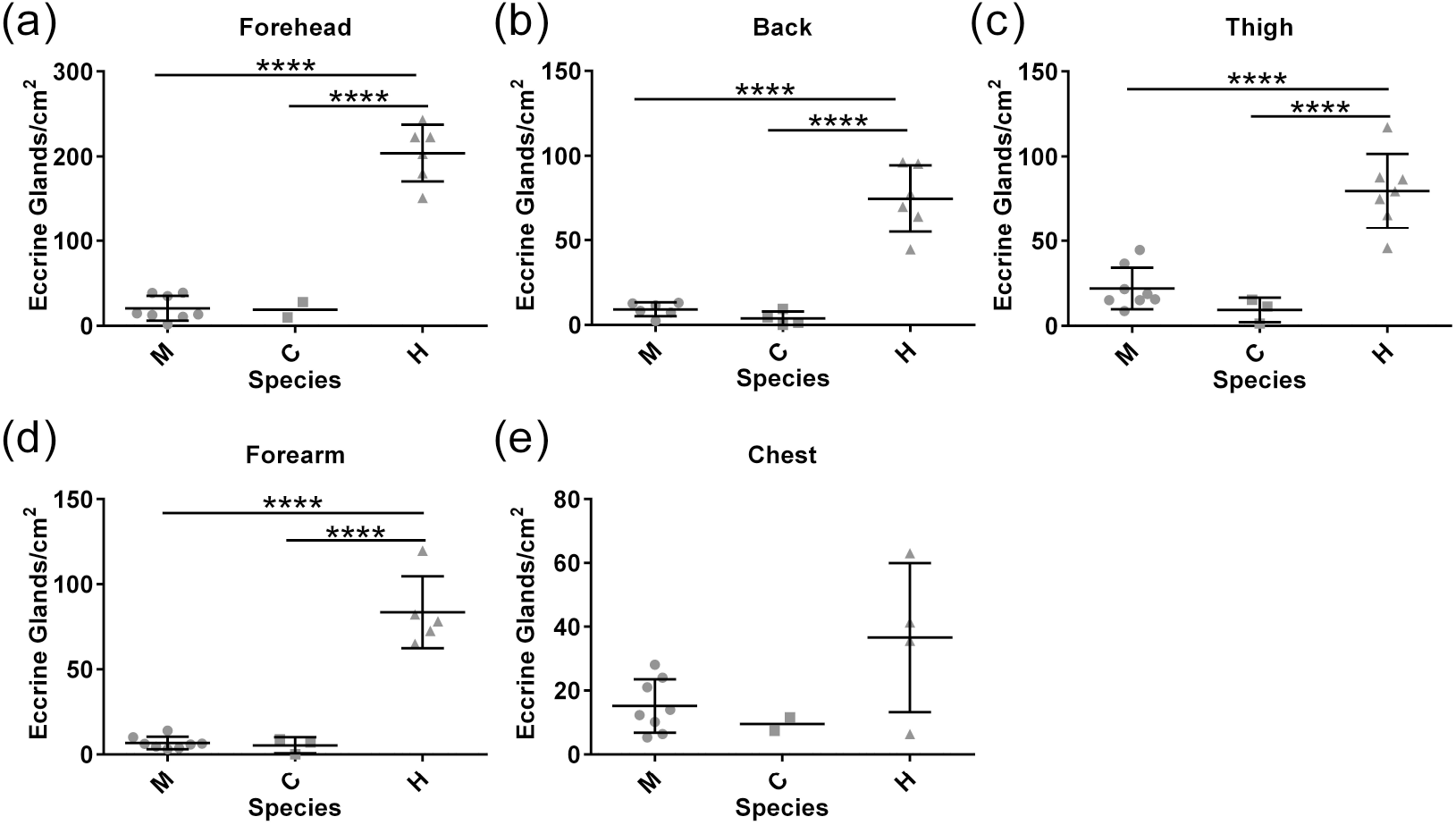
Eccrine gland density across body regions in three primates. a – e) Eccrine gland density is dramatically higher in humans than in macaques and chimpanzees, which are strikingly similar. Each point represents the average density of eccrine glands in biological replicates for each individual analyzed. Mean and standard deviation are shown for each species. Standard deviation not shown when sample size did not exceed two individuals. Abbreviations: macaque (M), chimpanzee (C) and human (H). Significance: * *P* < 0.05, ** *P* < 0.01, *** *P* < 0.001, **** *P* < 0.0001.

### Patterns of total ectodermal appendage density between species vary with body region

In most body regions, macaques had the greatest combined appendage density and chimpanzees the lowest (Fig. 4). In the back, chest and thigh, human and chimpanzee appendage density did not differ significantly (adjusted *P* > 0.05 for all comparisons), and in general was two to eight times lower than in macaques (Fig. 4b, c, e; adjusted *P* values: Back *P* = 0.0007 (M:C), *P* = 0.0153; Chest *P* = 0.1693 (M:C); *P* = 0.0359 (M:H); Thigh *P* = 0.0011 (M:C); *P* = 0.0231 (M:H)). In the forehead and forearm, however, total appendage density in humans was higher and more similar to macaques than chimpanzees (Fig. 4a, d; adjusted *P* values: *P* > 0.05 for H:M comparisons; Forehead *P* = 0.0232 (H:C); Forearm *P* = 0.0399 (H:C)). Total ectodermal appendage density was highest in the foreheads of macaques and humans, with no significant differences in combined appendage densities between the other sampled regions (SOM Fig. S3a, b). A similar trend was observed in chimpanzees (Fig. S3c).

**Fig. 4.**
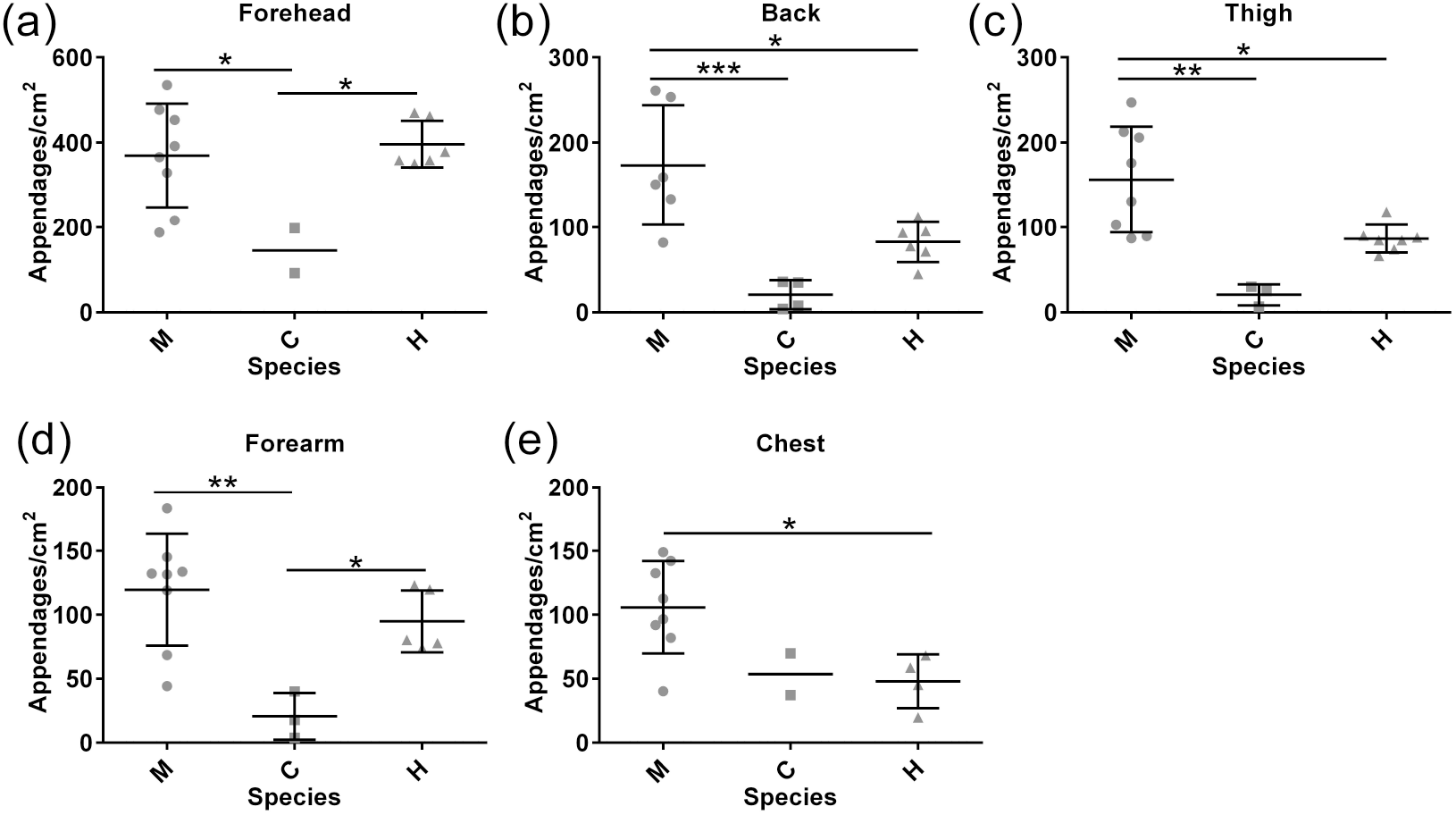
Appendage density across body regions in three primates. a – e) The total density of skin ectodermal appendages, based on combined hair follicles and eccrine glands, varies with body region. Each point represents the average density of appendages in biological replicates for each individual analyzed. Mean and standard deviation are shown for each species. Standard deviation not shown when sample size did not exceed two individuals. Abbreviations: macaque (M), chimpanzee (C) and human (H). Significance: * *P* < 0.05, ** *P* < 0.01, *** *P* < 0.001.

Because eccrine gland and hair follicle density have been shown to be inversely correlated in the inter-footpad skin of mice (Kamberov et al., 2015), we evaluated the relative abundance of each appendage type in each body region (SOM Fig. S4). We analyzed the *P* value and found no evidence for correlation between eccrine gland density and hair follicle density traits in any body regions (Spearman correlation test, *P* > 0.05 for all tested regions and SOM Fig. S4). Thus, differences in total appendage density between species seem to result from independent variation in the density of each appendage type within each body region.

## 4. Discussion

Our analysis of ectodermal appendages indicates that humans are indistinguishable in terms of hair follicle density from chimpanzees and that both species share reduced hair density compared to macaques. This finding suggests that the absence of visible fur cover in humans does not result from a loss of hair follicles per se, but instead favors a model of evolutionary changes in the morphogenesis of the individual hair follicles, which resulted in a shift from the large, pigmented terminal hair type characteristic of fur to the microscopic, unpigmented vellus hair type that has been described across most of the human body.

A relative decrease in body hair density between chimpanzees and macaques was also observed in the classical study of primate hair density by Schultz (1931) in which only hairs greater than two mm in length, thus a subset of body hairs, were analyzed (Schultz, 1931). In the same study, Schultz observed a similar trend when comparing hair density of Old World monkey species to that of great apes (Schultz, 1931). As previously suggested by Schwartz and Rosenblum (1981), one possible explanation for the relative decrease in hominid hair density is that hair density scales inversely with body surface area (Schwartz and Rosenblum, 1981). Though not working with human data, these authors argued that as thermal-generating volume increases to the power of 1.5 to body surface area from which heat can be lost, a reduction in insulating body hair would be advantageous to larger primates (Schwartz and Rosenblum, 1981). Extrapolating to humans, they and others (Newman, 1970; Montagna, 1972) proposed that hair density would have been similarly reduced in hominins and that the resulting reduction in protection from solar radiation would have favored the evolution of a mechanism of evaporative heat loss through sweating (Schwartz and Rosenblum, 1981). Despite their reduced fur cover, however, there is no evidence that chimpanzees employ a different mechanism of heat dissipation from macaques, nor one that is more similar to humans. If the reduction in hair density in the anthropoid lineage did indeed come at a cost of reduced protection from radiant heat, it is intriguing to speculate that this may have helped to influence the choice of habitat for these species.

Analysis of the relative eccrine gland densities revealed that compared to chimpanzees and macaques, human skin contains the greatest density of eccrine sweat glands in nearly all body regions surveyed with the possible exception of the chest, which was undersampled in this study. Our absolute measurements of density are in good agreement with the mean estimates made by Krause (1844) and the Keio lab as reported by Yas Kuno (1956) (Wagner, 1844; Kuno, 1956). Our measurements of the relative density of eccrine glands across body regions also accord with prior reports that the forehead/face region contains the greatest density of these appendages in the body proper (Wagner, 1844; Kuno, 1956; Szabo, 1962, 1967). In contrast to previous studies, however, we did not observe significant differences in the density of glands among other body regions, despite having analyzed more individuals and sampled larger areas from each body region than these prior studies (Wagner, 1844; Kuno, 1956; Ellis and Montagna, 1962; Szabo, 1962, 1967; Montagna and Yun, 1963; Montagna et al., 1964; Montagna, 1972; Johnson and Elizondo, 1974). One explanation for this difference may be a decrease in sampling error in our study, but we cannot rule out the possibility that this difference reflects individual variation. Resolving this difference will require examination of eccrine gland density in more individuals of all three species coupled with the development of a non-invasive method to score these organs. Even so, our data indicate that humans are indeed distinct from the other primates examined in having the highest eccrine gland density. These data support a model of eccrine gland expansion specific to the hominin lineage.

A recent analysis of eccrine gland traits in primates suggests that natural selection has acted to increase sweating capacity in primates living in arid and hot environments, linking the evolution of eccrine gland functionality to environmental selective pressures (Best and Kamilar, 2018). Taking our findings into account, it would be intriguing to evaluate eccrine gland density in a larger subset of primate species to determine if this trait may have been subject to similar evolutionary constraints.

Our data indicate that chimpanzees and macaques have essentially identical eccrine gland densities, which contradicts prior assessments in the same species (Montagna and Yun, 1963; Montagna et al., 1964; Montagna, 1972). The conclusion of these classical studies is that within catarrhines, eccrine gland density is lowest in Old World monkeys, intermediate in great apes, and reaches its zenith in humans (Montagna, 1963, 1972). These observations form the basis for current theory on the evolution of eccrine gland density within the catarrhines (Montagna, 1985; Folk and Semken, 1991). A source of the discordance between the results reported in this study and these earlier works may stem from the fact that earlier surveys provided qualitative assessments of the number of rather than the density of eccrine glands. Another possible source of difference may be the number of individuals analyzed. For example, while Montagna measured four chimpanzee individuals, as did we, he analyzed one macaque (Montagna et al., 1964) as compared to eight individuals in our study. In fact, no more than three individuals of any primate species were analyzed in any of these surveys. Finally, our data on eccrine gland density in macaques are in accord with the large-scale study of Elizondo (1988) which also found no regional differences in gland density (Elizondo, 1988). Smaller sample sizes, an absence of absolute eccrine gland numbers and what appear to be estimates rather than actual calculations of eccrine gland density call into question some of the conclusions drawn in the classical studies by Montagna and colleagues. More data are clearly needed, but our observation of equivalent eccrine gland density between chimpanzees and macaques is a more robust assessment of the relative phenotypes of these species. In light of these findings, we consider possible models to explain the evolution of a macaque-like density of these organs in chimpanzees.

The similarly low eccrine gland density of chimpanzees and macaques despite chimpanzees being almost an order of magnitude bigger raises the intriguing question of why, unlike body hair, was there no dramatic shift in the density of these glands between such differently sized primates? Experimental studies of non-human primates including macaques and chimpanzees indicate that sweating is activated during thermal stress in these animals and contributes to evaporative heat loss (Johnson and Elizondo, 1974, 1979; Hiley, 1976; Whitford, 1976; Elizondo, 1977, 1988; Mahoney, 1980; Kolka and Elizondo, 1983). While the design of these studies does not allow for determination of the relative contribution of apocrine versus eccrine glands to heat dissipation, the observation that eccrine glands are activated in this context strongly indicates that they perform a thermoregulatory function in these species. Consistent with a function in primate thermoregulation, a recent study found evidence of natural selection for increased eccrine functional capacity in primates living in hot and arid environments where thermal heat load would be exacerbated (Best and Kamilar, 2018). In the context of serving a thermoregulatory function, since chimpanzees are considerably larger than macaques, they have much less surface area to unit volume, and thus one would predict that selection would have acted to increase eccrine gland density in chimpanzees. Since this change did not occur, one possibility is that there was no selective benefit to increase eccrine density. Alternatively, the presence of thermally-activated apocrine glands in the skin of these primates buffered the need to evolve an increased eccrine gland density, a possibility supported by the observation that sweating patterns in chimpanzees are best correlated with regions of high apocrine gland number and sweating in this species occurs even in the presence of atropine, an inhibitor of muscarinic receptors whose activity is required for eccrine gland activation (Whitford, 1976). Alternatively, these species may have evolved other compensatory mechanisms directly related to the functionality of eccrine glands for heat dissipation, a view consistent with the report by Best and Kamilar (2018) that, compared to macaques, chimpanzees have increased eccrine gland glycogen content and capillarization, both traits associated with greater sweating capacity. Finally, an expansion of eccrine glands in chimpanzees may have been constrained by the benefit of maintaining a thick fur coat. While eccrine glands and hair follicles employ an overlapping set of genetic cascades to guide their development (Biggs and Mikkola, 2014; Cui and Schlessinger, 2015), a recent study in mice suggests that enhancement of the developmental program for eccrine gland formation can lead to defects in hair formation (Lu et al., 2016).

## 5. Conclusions

Our data are consistent with a model in which separable changes in the density of eccrine and hair follicle appendages occurred over the course of primate evolution. The reduced hair follicle density of hominids relative to macaques suggests that hair density was reduced in a common ancestor of chimpanzees and humans, after the split from Old World monkeys, perhaps as a result of shifting surface area to volume ratios in larger anthropoids (Schwartz and Rosenblum, 1981; Sandel, 2013). In contrast, humans significantly diverge from the ape and macaque condition in terms of eccrine gland density. In support of a model of human-specific increase in eccrine gland density, we find no evident correlation between eccrine gland and hair follicle density in any of the primates studied. From a developmental perspective this independence between eccrine and follicle density would be facilitated by the fact that hair follicle specification occurs prior to the onset of eccrine gland formation during human gestation (Szabo, 1967; Lu et al., 2016).

Establishing the definitive hair and eccrine gland density traits of humans and our closely related primate relatives will not only help elucidate the genetic changes that drove their emergence, but will also hone current anthropological theory on the evolutionary history of our lineage and the selective pressures that drove adaptive changes in this context.

## Acknowledgments

This work was supported by the National Institutes of Health [grant numbers R01HD032443, 1995 to CJT; R21 AR066289, 2014 to BAM; 1DP2OD006514–01, 2010 to PCS], a grant from the Harvard University Science and Engineering Committee Seed Fund for Interdisciplinary Science to CJT, DEL, PCS and BAM, funding from the American School of Prehistoric Research to DEL, and a Packard Foundation Fellowship in Science to PCS. The authors acknowledge the Yerkes National Primate Research Center (NIH P51 RR000165, 2011), the Harvard Museum of Comparative Zoology, the New England Primate Research Center (NIH P51 RR000168, 2011), and the Anatomical Gift Program at Harvard Medical School for the tissue provided for this study. Declarations of interest: none.

## Abbreviation

acr: acrosyringium
sc: eccrine gland secretory coil
hf: hair follicle
sb: sebaceous gland
ap: arrector pili muscle

